# CHANging Consciousness Epistemically (CHANCE): An empirical method to convert the subjective content of consciousness into scientific data

**DOI:** 10.1101/495523

**Authors:** Daisuke H. Tanaka, Tsutomu Tanabe

**Affiliations:** Department of Pharmacology and Neurobiology, Graduate School of Medicine, Tokyo Medical and Dental University (TMDU), 1-5-45 Yushima, Bunkyo-ku, Tokyo 113-8519 Japan

## Abstract

The content of consciousness (cC) constitutes an essential part of human life and is at the very heart of the hard problem of consciousness. The cC of a person (e.g., study participant) has been examined *indirectly* by evaluating the person’s behavioral reports, bodily signs, or neural signals. However, the measures do not reflect the full spectrum of the person’s cC. In this paper, we define a method, called “CHANging Consciousness Epistemically” (CHANCE), to consciously experience a cC that would be identical to that experienced by another person, and thus *directly* know the entire spectrum of the other’s cC. In addition, the ontologically subjective knowledge about a person’s cC may be considered epistemically objective and scientific data. The CHANCE method comprises two empirical steps: (1) identifying the minimally sufficient, content-specific neural correlates of consciousness (mscNCC) and (2) reproducing a specific mscNCC in different brains.

## Introduction

A hungry person on eating an apple may consciously experience pleasant feelings. A person who is hurt may consciously experience pain. These subjective conscious experiences constitute a core part of human life and are central to accurately understanding the nature of consciousness (Chalmers, 1995; M. Tye, 2018). This conscious experience is called “content of consciousness” (cC) [Koch, Massimini, Boly, and Tononi, 2016], and it appears to be similar to other commonly used terms such as “phenomenal consciousness,” *“what it is like* aspect of experience,” or “qualia.” In this paper, the term “cC” is used synonymously with the aforementioned terms.

The cC exists only as experienced by each individual and is thus ontologically subjective, while the brain exists without the experiences of individuals and is thus ontologically objective (Searle, 1998). This fact raises an intriguing question, called the hard problem of consciousness (Chalmers, 1996): “How does the subjective cC arise from an objective brain?” Numerous scientific studies have been conducted in experimental psychology and cognitive neuroscience to reveal the neural basis of cC, yielding significant insights (Crick and Koch, 1990; Craig, 2009; Dehaene and Changeux, 2011; Freeman, 2007; Koch, 2004; Koch et al., 2016; Lau and Rosenthal, 2011; Tsuchiya, Wilke, Frassle, and Lamme, 2015). Using typical experimental paradigms, researchers record and compare the elicited neural activity, based on whether individuals (e.g., study participants) experienced a specific cC. A participant’s cC is examined *indirectly* through a verbal report or by pressing of a button in response to a “yes” or “no” question such as “Did you see a dot?” (refer to Figure 1a, thin arrow) [Del Cul, Baillet, and Dehaene, 2007; Lutz, Lachaux, Martinerie, and Varela, 2002; Ress, Backus, and Heeger, 2000; Sandberg, Timmermans, Overgaard, and Cleeremans, 2010; Super, Spekreijse, and Lamme, 2001; Tong, Meng, and Blake, 2006]. However, both of these reports (or more generally, behavioral reports) about a person’s cC vary because of a shift in the criterion with regard to what constitutes a “yes*”* response (i.e., having a cC) for a specific person, particularly when the cC is at or near perceptual thresholds (Kunimoto, Miller, and Pashler, 2001). Some researchers supplement these reports with confidence measures such as the Perceptual Awareness Scale (Sandberg et al., 2010) in which responses can range from “no experience” to “absolutely clear image” or a post-decision wagering (e.g., “Would you bet that your response was correct?” [Persaud, McLeod, and Cowey, 2007]) to gain additional information about a person’s cC (Kunimoto et al., 2001; Schurger and Sher, 2008). However, these confidence measures do not always match the person’s behavioral reports about the cC (Kanai, Walsh, and Tseng, 2010). Hence, the behavioral measures of cC remain debatable (Dehaene and Changeux, 2011).

**Figure 1:**
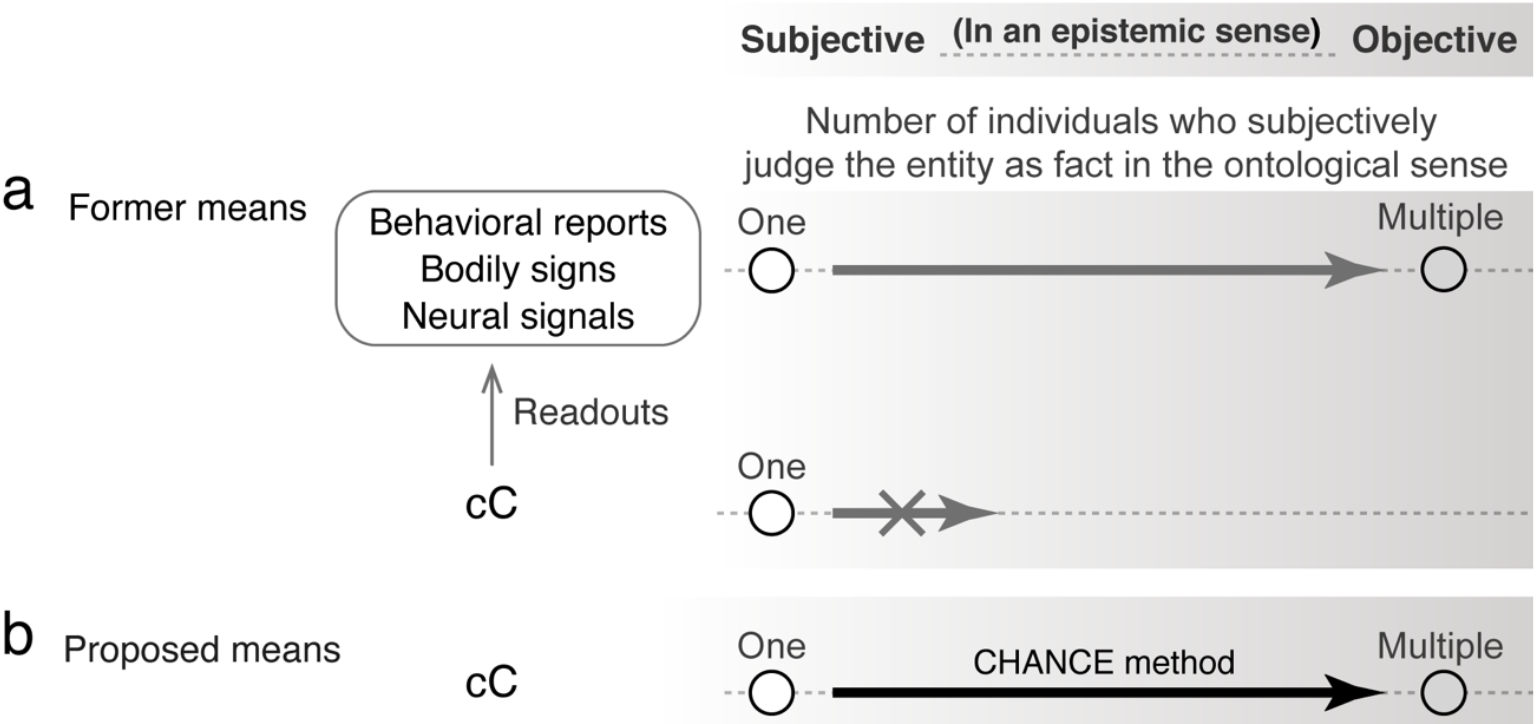
The CHANCE method converts a cC from being epistemically subjective to being epistemically objective. (**a**) The former conventional scientific means of addressing cC is depicted. The cC is judged as a fact only by the original person and it is impossible to objectify it epistemically (bold lower arrow with the “X”). Behavioral reports, bodily signs, or neural signals are “readouts” of the cC (thin arrow). The readout data are subjectively judged as facts in the ontological sense by multiple relevant individuals (e.g., experts in the research field); thus, they are considered epistemically objective and scientific data (thick upper arrow). However, no readout fully reflects the cC. (**b**) A proposed means of addressing the cC is illustrated. The cC is empirically changed from being epistemically subjective to epistemically objective. If the cC is subjectively judged as fact in the ontological sense by multiple relevant individuals, then it would be considered epistemically objective and therefore, scientific data (thick arrow)

In addition, behavioral reports involve various cognitive functions such as attention (Koch and Tsuchiya, 2007; Lamme, 2003), working memory (Soto and Silvanto, 2014), expectation (Kok, Rahnev, Jehee, Lau, and de Lange, 2012; Melloni, Schwiedrzik, Muller, Rodriguez, and Singer, 2011), and meta-cognition (Kanai et al., 2010). If cC is experienced in the absence of these cognitive functions, the participant will not be able to report the cC, and thus, researchers may underestimate the putative neural activity underlying the cC. In addition, the neural activity underlying these cognitive functions is difficult to discriminate from those underlying cC (Cohen and Dennett, 2011; Koch et al., 2016; Tsuchiya et al., 2015), causing an overestimation of the putative neural activity underlying the cC. Collectively, behavioral measures do not entirely reflect a person’s cC and can cause both an underestimation and overestimation of the neural activity underlying the cC.

Several studies have assessed cC through bodily signs, such as pupil size (Frassle, Sommer, Jansen, Naber, and Einhauser, 2014), or through neural signals in the absence of behavioral reports (Garcia, Srinivasan, and Serences, 2013; Haynes, 2009; Horikawa, Tamaki, Miyawaki, and Kamitani, 2013; Nishimoto, Vu, Naselaris, Benjamini, Yu, and Gallant, 2011) [see Figure 1a, thin arrow]. These approaches may overcome some of the aforementioned problems in the report-based paradigm. However, they may also cause both an underestimation in the putative neural activity underlying the cC by missing percepts due to a no-report and an overestimation by including unconscious neural processing (Tsuchiya et al., 2015).

Furthermore, current methods, regardless of whether they are based on reports or no-reports, are limited to evaluating a person’s responses to a simple question (e.g., “Did you see a dot?”) or to a simple stimulation (e.g., viewing a flower picture), and consequently, only provide limited cC information. No behavioral report, bodily sign, or neural signal reflects the entire spectrum of a person’s cC (Chalmers, 1996, 1999; Nagel, 1974; Velmans, 2007) [Figure 1a, open arrow]. Therefore, it is crucial for researchers to develop a novel method that can be used to accurately know a person’s cC.

An ideal method for researchers to accurately know the person’s cC is to consciously experience an identical cC. However, this idea raises both technical and philosophical questions: Can a cC be induced in a researcher such that it is identical to the participant’s cC? Can the researcher’s cC be considered scientific data? To address these questions and define a novel method to accurately know another person’s cC, we first evaluated the sufficient conditions to be considered scientific data. We then defined a method, called “CHANging Consciousness Epistemically” (CHANCE), that converts subjective cC to scientific data. This method would induce the participant’s cC in a researcher, and the researcher’s cC that is identical to the participant’s cC would provide scientific data.

One may argue that we propose a novel method of evaluating a behavioral report, bodily sign, or neural signal with the guise of being *direct.* However, this is not the case, as we propose that the CHANCE method allows one to experience and *directly* know the cC of other individuals without using those measures. The CHANCE method does not involve any of the measures of a person’s cC but provides researchers with the conscious experience and knowledge regarding another person’s cC.

## The Content of Consciousness Can Become Scientific Data in Theory

### Scientific Data are Epistemically Objective Data and Vice Versa

Scientific data are those that may be known to be true or false in a way that does not depend on the preferences, attitudes, or prejudices of individuals (Chalmers, 1996, 1999; Descartes, 1644/1972; Galileo, 1623/1957; Searle, 1998; Velmans, 2007); thus, scientific data are epistemically objective data and vice versa (Searle, 1998). Therefore, if a cC was epistemically objective, it would be considered scientific data. It is widely believed, however, that the cC of an individual cannot be known to be true or false in a way that does not depend on the preferences, attitudes, or prejudices of individuals; thus, the cC is not epistemically objective but subjective. This is indeed the reason for researchers to use behavioral reports, bodily signs, or neural signals as “readouts” to know a person’s cC in scientific investigations (Figure 1a).

### The Content of Consciousness Can Become Epistemically Objective

Epistemically objective and subjective entities have been considered qualitatively different (Berridge, 1999; Berridge and Kringelbach, 2015; LeDoux, 2014; K.M. Tye, 2018). However, the subjective–objective distinction seems more blurred than what many have previously acknowledged. For example, researchers in one scientific laboratory may repeatedly conduct experiments to obtain data, whereas a specific researcher in another laboratory may conduct the same experiment only once. Most researchers would hopefully agree that, although the data obtained in each laboratory would be objective, the data obtained in the first situation would be more faithful to the facts (i.e., the truth). Therefore, this information would be more objective than that in the second scenario. This greater objectivity is due to the fact that, in the latter situation, data may be obtained by chance or because of a specific researcher’s subjective biases (i.e., personal beliefs or preferences). Thus, the epistemic objectivity of a datum (i.e., an entity) may exist in degrees (Reiss and Sprenger, 2017). In the epistemic sense, the terms “subjective” and “objective” may be at the opposite poles of the same axis, and most entities between these polarities have some degree of objectivity. That is, objective and subjective entities are not qualitatively different and the border between them may not exist. This argument raises the possibility that the cC which has been believed to be absolutely subjective may have a certain degree of epistemic objectivity (Figure 1b).

Three features appear to be involved in determining an entity’s degree of epistemic objectivity. Firstly, it is reasonably assessed by individuals who have the ability to judge how faithful the entity is to fact (Reiss and Sprenger, 2017). For example, the faithfulness of scientific results is usually judged by scientists in relevant research fields (e.g., editors and reviewers of journals). Secondly, each individual’s judgment is always achieved subjectively in the ontological sense (Vaerla, 1996; Velmans, 1999). When a scientist observes experimental results or scientific data and judges its faithfulness to fact, it is a conscious and subjective effort. Lastly, a large number of individuals judging the entity as fact results in greater faithfulness, and consequently, greater epistemic objectivity. This argument is consistent with the “intersubjective agreement” in which a consensus among different individual judgments often indicates objectivity (Steup, 2018). Collectively, a specific entity including a cC would be epistemically objective if multiple relevant individuals subjectively judged it as fact in the ontological sense (Figure 1b).

## A Method to Render the Content of Consciousness as Scientific Data

CHANging Consciousness Epistemically (CHANCE) enables a specific cC to be subjectively judged as a fact in the ontological sense by multiple relevant individuals, and thus undergoes a change from being epistemically subjective to epistemically objective and therefore can be considered scientific data (Figure 1b). The CHANCE method consists of two empirical steps: (1) identifying the minimally sufficient, content-specific neural correlates of consciousness (mscNCC) and (2) reproducing a specific mscNCC in different brains.

### Step One: Identifying Neural Bases

Specific neural bases in the human brain are sufficient to produce cC (Crick and Koch, 1990; Craig, 2009; Dehaene and Changeux, 2011; Freeman, 2007; Koch, 2004; Koch et al., 2016; Lau and Rosenthal, 2011; Tononi and Koch, 2015). Koch et al. (2016, p. 308) argued that “the neurons (or, more generally, neuronal mechanisms), the activity of which determines a particular phenomenal distinction within an experience” are the content-specific neural correlates of consciousness (NCC). Chalmers (2000, p. 31) defines an NCC for a cC as follows: “An NCC (for content) is a minimal neural representational system *N* such that representation of content in *N* is sufficient, under condition *C*, for the representation of that content in consciousness.” Inspired by these concepts, we assumed that there exists a neural event that is minimally sufficient to produce a specific cC without any other support mechanism. We named the event mscNCC [Figure 2a, Step 1]. When an mscNCC occurs in a person’s brain, a specific cC should be experienced in all possible instances and conditions; however, even without the mscNCC, the person may still experience the cC through neural events other than the mscNCC. An mscNCC is self-sufficient to produce a specific cC without any other support mechanism. This appears to contrast with Chalmers’ NCC (for content) [Chalmers, 2000, pp. 25–26] which asserts that “nobody (or almost nobody) holds that if one excises the entire inferior temporal cortex or intralaminar nucleus and puts it in a jar, and puts the system into a relevant state, it will be accompanied by the corresponding state of consciousness.” We claim that if an mscNCC that produces a specific cC were isolated from the human brain and placed in a jar, it would still produce the cC. An mscNCC alone is essentially truly sufficient to produce a specific cC in all possible instances and conditions. An mscNCC produces only one specific cC. To ensure that an mscNCC is minimal, each neuronal, synaptic, and molecular event — or more generally, a neural event comprising the mscNCC — should be tested to determine whether it is indeed necessary to produce the specific cC.

**Figure 2:**
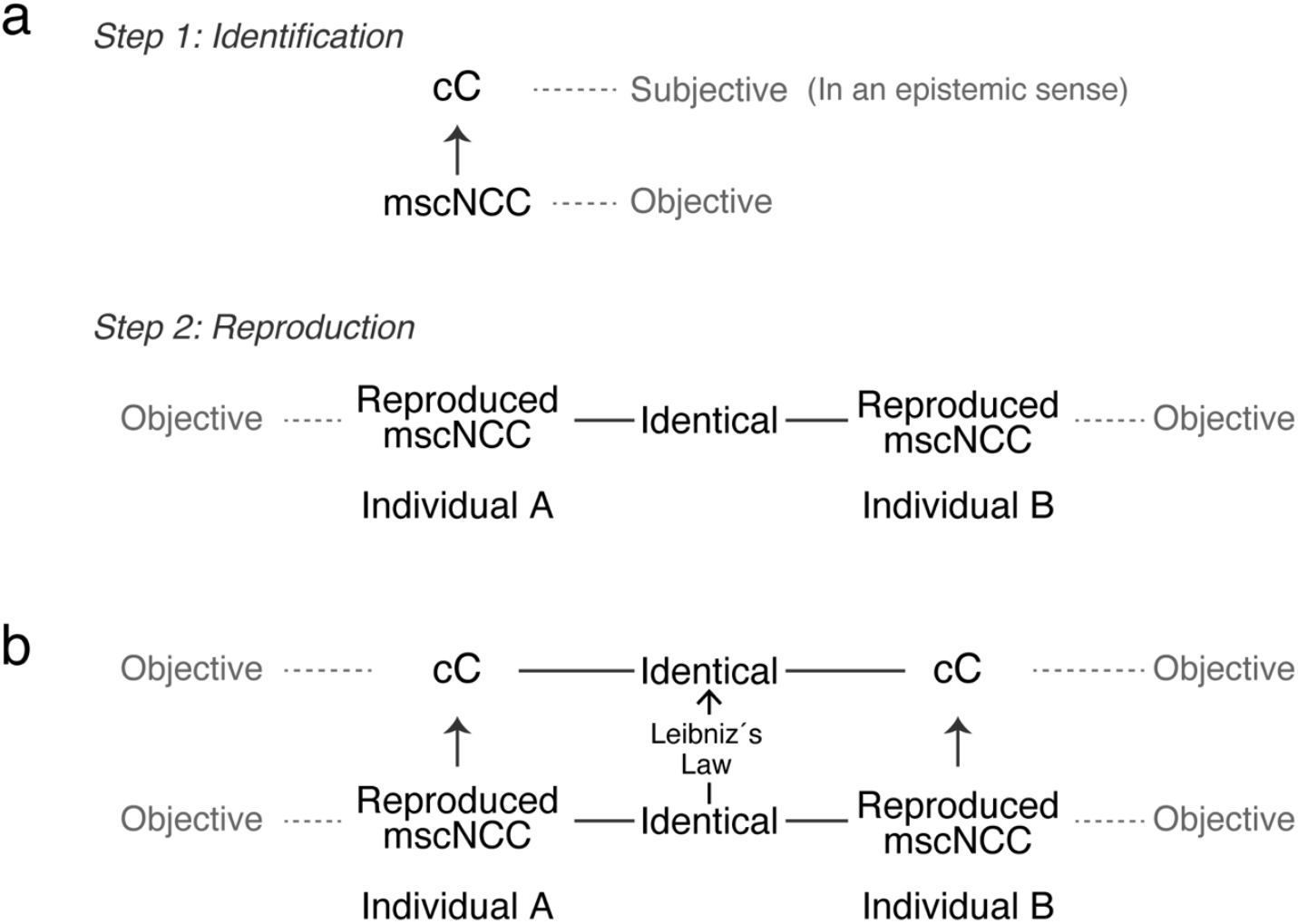
The CHANCE method and a logical consequence of its creation. (**a**) The two steps in CHANCE are depicted. Step 1 involves identifying an mscNCC, defined as the minimum neural events that are sufficient to produce only one specific cC. An mscNCC itself is epistemically objective; however, a cC is epistemically subjective, making the experiments and the results obtained in this step nonscientific. Step 2 involves reproducing an mscNCC in different brains. The reproduced mscNCCs among different brains are identical. This step contains only the epistemically objective entity, the mscNCC; thus, experiments and the results obtained in this step are scientific. (**b**) A logical consequence of the verification of the two steps in CHANCE (a). The mscNCC that produces a specific cC in Individual A is identical to the mscNCC of Individual B. The occurrence of the mscNCC in Individual B should produce an identical cC to that of Individual A as a logical consequence of Step 1, Step 2 (a), and Leibniz’s Law. An identical cC would be subjectively judged as a fact in the ontological sense by multiple individuals (i.e., Individuals A and B). If relevant individuals capable of judging the objectivity of the cC joined the experiment as participants, the shared identical cC would be considered epistemically objective, thus providing scientific data.

One may argue that a few consciousness researchers, except for the proponents of panpsychism (Koch et al., 2016; Tononi and Koch, 2015), would assume that an mscNCC can still produce a cC if it is isolated from the human brain. This argument may originate from intuition or common sense. Many consciousness researchers would likely agree that, if a whole human brain was placed in a jar and activated appropriately, the brain would produce a cC. In this condition, not all neural events in the brain would be necessary to produce the cC; hence, the unnecessary neural events could be removed from the brain. By repeated removals, only the mscNCC would ultimately remain in the jar and still produce a cC. Therefore, it is not unrealistic to assume that an mscNCC in a jar produces a cC.

To empirically identify an mscNCC, the relevant neural events need to be empirically induced with high spatiotemporal resolution, whereas the effects of the induction on a cC need to be consciously and subjectively experienced by a researcher or individual who intends to evaluate the effects. Thus, the brain of a researcher or individual who intends to evaluate the results needs to be empirically manipulated. The results obtained by the experiment would be a cC and only available to the researcher or individual whose brain was manipulated. Therefore, those results would be epistemically subjective (Figure 2a, Step 1). This epistemically subjective result would make the experiment nonscientific. However, this methodological limitation would not decrease the confidence obtained by each participant who evaluates cC-containing results, compared to standard scientific results, because both methods would provide ontologically subjective knowledge and confidence to each individual. The relevant neural events would be viewed as an mscNCC, if the following conditions were verified: (1) a researcher or individual whose brain is manipulated experiences only one specific cC, when the relevant neural events are induced (i.e., verification of *sufficiency)* and (2) a researcher or individual whose brain is manipulated does not experience the specific cC when any neural event among the relevant ones is inhibited, even if all other neural events among the relevant ones are induced (i.e., verification of *minimality).* The manipulated individual should experience and know a specific cC, when a specific mscNCC occurs, regardless of whether *any other neural events* occur. Once an appropriate mscNCC is identified, the occurrence of the mscNCC would indicate the production of a specific cC.

One may argue that it is unrealistic to attempt to verify the two aforementioned conditions for identifying an mscNCC. Indeed, the neural events that are crucial in sustaining life, such as the neural events controlling respiration, may need to be inhibited temporarily to test whether they are included in the mscNCC. For nonhuman animals, several interesting techniques have been developed to manipulate neural activities, such as combining optogenetics with modern methods in system neuroscience (Kim, Adhikari, and Deisseroth, 2017). However, the spatiotemporal precision of the current techniques appears insufficient to conduct the experiments necessary to verify both criteria. These are technical difficulties, rather than theoretical limitations, and may be overcome in the future.

One may also argue that it is implausible to assume that an mscNCC produces only one specific cC, and no other cC, because cCs are highly sensitive to context. For example, the brightness of two patches with identical absolute luminance is experienced differently when the patches are surrounded by different contexts (Adelson, 2000). However, this situation does not necessarily mean that a specific mscNCC produces two different cCs, depending on other neural activities. This situation is instead interpreted as follows: the brightness of patch A surrounded by context A is produced by a specific mscNCC, whereas the brightness of patch A surrounded by context B is produced by a different mscNCC. That is, the different brightness of identical patches in absolute luminance surrounded by different contexts is produced by different mscNCCs. Alternatively, specific stimulus information (e.g., the absolute luminance of a patch) induces a specific mscNCC in a specific situation but induces another mscNCC in a different situation, depending on other information (e.g., the surrounding context of the patch).

Nevertheless, some researchers may argue that the requirement of an mscNCC to establish the CHANCE method results in a circular argument: Establishing CHANCE may enable a cC to be considered as scientific data and lead to classification by neural bases. However, to establish CHANCE, one first needs to know what these bases are. This argument results from a lack of distinction between the degree of epistemic objectivity with regard to the cC before and after the establishment of CHANCE. When using the CHANCE method, a cC is studied in an epistemically subjective (i.e., nonscientific) manner during Step 1 (Figure 2a); however, when both Step 1 and 2 in CHANCE are verified, a cC is studied in an epistemically objective (i.e., scientific) manner (Figure 2b). Thus, although epistemically subjective knowledge regarding the neural mechanism of a cC is used to establish CHANCE (Figure 2a, Step 1), when CHANCE is established, the epistemically subjective knowledge can then be converted to epistemically objective scientific knowledge (Figures 1b and 2b). Ontologically subjective cC becomes epistemically objective; thus, it would be considered scientific data (Figure 2b).

### Step Two: Reproducing the Neural Bases in Different Brains

Next, a specific mscNCC is reproduced in different brains (Figure 2a, Step 2). To achieve this, sophisticated technologies need to be developed. For example, if the essential neural events of the mscNCC were specific activities in specific neural networks such as those in the Global Neuronal Workspace (GNW) [Baars, 1989; Dehaene and Changeux, 2011; Dehaene, Kerszberg, and Changeux, 1998], the same patterns of activation should be reproduced. The mscNCCs reproduced in different brains should be identical (Figure 2a, Step 2). To ensure the identicalness, the precise identification of the neural events of the mscNCC — for example, specific neural or synaptic activity patterns — in the aforementioned Step 1 is crucial. Recent developments in noninvasive human brain-to-brain interface (Lee, Kim, Kim, Lee, Chung, Kim, and Yoo, 2017; Mashat, Li, and Zhang, 2017; Yoo, Kim, Filandrianos, Taghados, and Park, 2013) may aid in reproducing some neural events in different brains. However, current precision tools seem inadequate for reproducing potential neural events of an mscNCC such as GNW activity. Therefore, technical developments are needed to achieve this step.

### Verification of the Two Steps Makes the Content of Consciousness Epistemically Objective

If the previous two steps are verified, then the occurrence of a specific mscNCC would produce a specific cC (Figure 2a, Step 1), and a specific mscNCC would be reproduced in different brains (Figure 2a, Step 2). Based on Leibniz’s Law which states “that for anything *x* and for anything y, if *x* is identical with *y* then *x* and *y* share *all* the same properties” (M. Tye, 2018, p. 11), the reproduced identical mscNCCs should share *all* of the same properties, including the ability to produce identical cCs in different individuals (Figure 2b). The relevant individuals who judge the faithfulness of the cC can then join the experiment. The identical cC that is shared and judged subjectively as a fact in the ontological sense by multiple relevant individuals can be considered epistemically objective (Figures 1b and 2b). Velmans accordingly argued that shared experiences among multiple individuals might be public and objective; “to the extent that an experience … can be *generally* shared (by a community of observers), it can form part of the database of a communal science” (1999, p. 304).

One may posit that it is difficult to ascertain that a cC in multiple individuals does not vary according to the influence of the surrounding unreproduced neural activity. This argument appears to arise from a misunderstanding at Step 1, which focuses on the mscNCC that produces only one specific cC, regardless of the activity of any other surrounding neurons (Figure 2a, Step 1). Even if the surrounding unreproduced neural activity varied among individuals, these neural activities would not influence the cC produced by mscNCC because a specific cC can be entirely produced solely by a specific mscNCC under any other neural activity (Figure 2a, Step 1).

Some readers may suggest the need to demonstrate that the cC shared among multiple individuals is indeed identical. As previously mentioned, the equivalence of the cCs experienced and known by each individual is a logical consequence of the two steps in CHANCE and Leibniz’s Law — namely, a specific mscNCC produces a specific cC, regardless of any other neural activity (i.e., Step 1), and an identical mscNCC is reproduced in multiple individuals (i.e., Step 2); thus, identical mscNCCs should produce identical cCs. Therefore, the identicalness of shared cCs among multiple individuals is logically plausible without the direct empirical demonstration of the equality.

One may argue that, in the scenario of an *inverted spectrum* (Block, 1980, 1990; Shoemaker, 1982), an mscNCC that produces red content in one individual can be identical to an mscNCC that produces green content in another individual. This argument can originate from misunderstandings in Step 1 and Leibniz’s Law: if a specific mscNCC produced a specific cC regardless of any other activities (i.e., Step 1), then the identical mscNCCs reproduced in different brains should produce an identical cC (i.e., the logic of Leibniz’s Law). Therefore, if the mscNCCs reproduced in two individuals are identical, and if an mscNCC in one individual produces red content, another identical mscNCC in another individual should produce red content, not green.

## Discussion

### Requirements to be considered scientific data should be defined

If the degree of the epistemic objectivity of an entity has been reasonably judged by relevant individuals (Reiss and Sprenger, 2017), it remains unclear as to who would judge the degree of epistemic objectivity of shared identical cCs (Figure 2b). It also remains unclear how many relevant individuals are necessary to judge a cC as a fact and what degree of epistemic objectivity is essential for a cC to be considered scientific data. We argue that it is essential to develop a standard to quantify the degree of epistemic objectivity of specific entities, as well as on the same, to be considered scientific data.

### An Answer to Nagel’s Question and the Denial of the “Philosophical Zombie”

If an identical cC were shared among multiple individuals (Figure 2b), scientists would be able to respond to Nagel’s (1974) well-known philosophical question: “What is it like to be a bat?” The question indicates that “to know whether you, the reader, are conscious, I must know what it is like to be you” (Baars, 1996). This request implies that an observer (e.g., a researcher) should somehow share the cC of a subject (Baars, 1996), which would be achieved upon establishing CHANCE (Figure 2b). The researcher would share an identical cC with the participant and subsequently have “observer empathy” (Baars, 1996), knowing what it is like to be the other person. Thus, the researcher would know that the participant does not experience the *inverted spectrum* (Block, 1980, 1990; Shoemaker, 1982) and that the individual is not a *philosophical zombie* behaving normally without cCs (Chalmers, 1996).

### Addressing Obstacles in First-Person Data

First-person data concerning the cC contain something that is excluded in heterophenomenology (Dennett, 1991, 2001) and in critical phenomenology (Velmans, 2007) but is centrally important to the nature of the cC (Chalmers, 2013). Chalmers claims that first-person data are accompanied by obstacles when they are used in the science of consciousness. He claims that “first-person data concerning subjective experiences are directly available only to the subject having those experiences” (p. 32) and only *indirectly* available to others through their readouts (Figures 1a). However, if a person’s cC is shared among others (Figure 2b), the first-person data concerning the cC would be *directly* available to them, making the first-person data concerning the cC nonexclusive. Chalmers also claims that current “methods for gathering first-person data are quite primitive” (p. 33). If a person’s cC is shared among others, then gathering first-person data would be unnecessary because the first-person data concerning the cC would be *directly* available to others (Figures 1b and 2b). Chalmers contends that the general formalism to express first-person data is lacking, but is necessary for data gathering and theory construction. Contrastingly, gathering first-person data would be unnecessary if a person’s cC is shared, thereby removing the need for formalism. However, the development of formalism would be necessary to record, in writing, the results of experiments and to construct and describe a theory explaining the relationship between a cC and its underlying neural mechanisms. Therefore, epistemic objectification of a cC would overcome several, if not all, obstacles involving first-person data (Chalmers, 2013), and would introduce a new method to incorporate them into the science of consciousness.

## Acknowledgements

We thank Dr. F. Murakami and Dr. I. Fujita at Osaka University and Mr. R. Matsumura and Mr. S. Inaba at TMDU for their helpful comments and discussions. This work was supported by JSPS KAKENHI Grant Numbers JP26890011, JP16K07024 and JP19K06938, Takeda Science Foundation and TOBE MAKI Scholarship Foundation to D.H.T.

## Citation information

The current paper is formally accepted in The Journal of Mind and Behavior. If you need to cite the current paper, then please use the following information:

> Tanaka DH and Tanabe T. CHANging Consciousness Epistemically (CHANCE): An empirical method to convert the subjective content of consciousness into scientific data, *J. Mind Behav*. 40 (3/4), 177–190 (2019)

